# Innate antiviral systems are major defensome components that drive prophage distribution in *Acinetobacter baumannii*

**DOI:** 10.1101/2024.10.26.620419

**Authors:** Antonio Moreno-Rodríguez, Alejandro Rubio, Andrés Garzón, Younes Smani, Antonio J. Pérez-Pulido

## Abstract

Phages are guilty of killing daily almost half of bacterial cells, while bacteria have developed defense mechanisms that number in the dozens. Individual defense systems are gained and lost by genomes of the same species, depending on their fitness advantage. Thus, some genomes have a certain combination of defense systems, while other genomes act as a reservoir for the rest of the systems, thus constituting the so-called pan-immune system of the species. Here we have analyzed thousands of genomes of the bacterium *Acinetobacter baumannii*, an opportunistic pathogen of humans of great clinical concern, and we have found 81 different defense systems heterogeneously distributed. By analyzing how these systems combine, we have found that more than half of the genomes lack the universal DNA-methylating restriction-modification systems (R-M) and harbor an alternative innate SspBCDE system that performs a DNA phosphorothioate modification. In addition, the adaptive CRISPR-Cas systems could act synergistically to the R-M systems, based on their frequency of co-appearance. The presence of one or the other innate system could modulate the evolution of the genomes of this species, causing them to present a different profile of phages integrated into the bacterial genome. We have also observed that the presence of many defense systems is associated with the presence of a higher number of prophages, which could be due to the fact that the prophage carries the system, or that the bacterium would not need these systems in environments where the phage is absent.

## Introduction

Bacteriophages, also known as phages, are bacteria-specific viruses that outnumber bacteria by a ratio of 10 to 1 (1). Worldwide, it is estimated that 10^24^ bacterial infections by phages occur every second, which would eliminate 5-50% of bacteria in environments such as the oceans every day (2).

Phages usually attach to bacteria through surface structures such as *pili* or *flagella* and, using membrane proteins as specific receptors, inject their genome into the bacterial cell (3). As a result, they are often highly specialized, and a specific phage usually only recognizes bacteria of a certain species or even only strains that have a particular surface protein involved in the attachment phase or entry of the genome into the cell.

Bacteria defend themselves against phages using different systems. First, they present mechanisms that block the adsorption of the phage into the bacterium, thus preventing the entry of its genome (4). Once the phage genome enters the bacterium, there are three main defense systems, which have been known for decades. By far the most studied are the innate restriction-methylation (R-M) systems that appear in 74% of bacterial genomes (5) with no taxonomic group of bacteria or archaea lacking them entirely (6). They are based on prior labeling of the bacterial genome with methyl groups to subsequently destroy the unlabeled genome of possible viruses entering the bacterium by means of site-specific endonucleases (7). On the other hand, CRISPR-Cas systems are adaptive immunity systems, discovered in the 2000s (8, 9) and appearing in 39% of bacterial genomes and 90% of archaea (10). These act similarly to a vaccine, as they can store fragments of phage genomes from previous infections called spacers, which allows them to act more quickly and efficiently against future infections of the same phage. Spacer formation appears to be enhanced in cells that also carry R-M systems, as these limit phage replication through phage DNA degradation, and cleavage products can be captured by the Cas adaptation machinery (11, 12).

Finally, abortive infection (abi) systems function as a programmed cell death induction mechanism triggered by phage infection, preventing the spread of the phage within the bacterial population (13). An example of these are the CBASS systems, which in response to viral infection and based on a signaling process, trigger cell death (14). Another type of abi-associated systems are toxin-antitoxin systems, which add to the arsenal of bacterial defense systems in genomes carrying the two components of these type of systems (15).

All these antiviral systems are unevenly spread in bacteria, and even within the same species they may appear in some genomes and not in others (16). Thus, they are part of the accessory genome of the species, which supports their dispersal by horizontal transfer (17). In fact, in recent years many more anti-phage defense systems have been discovered, all of which means that we can define the pan-immune system or defensome of a species as the set of defense systems that it can present in their genomes (18, 19).

R-M are a kind of innate systems in bacteria that defend them from virtually any phage lacking the system’s methylation pattern. Uncommonly, there are bacteria that lack them and instead have other innate systems based on a different DNA labeling pattern, such as Ssp, BREX and Dnd (20, 21). SspABCD system, a subtype of the Ssp systems, is among the most studied (22, 23). They label the resident DNA by modifying the sugar-phosphate backbone of it, replacing the non-bridging oxygen with sulfur in a phosphorothioation reaction. Other Ssp subtype, SspBCDE, is known to confer high protection in *Escherichia coli* against coliphages (24).

All these systems are usually found in genomic islands and archipelagos, integrated in hotspots, which allows for easier discovery (25, 26). Based on this particularity, 29 groups of antiviral genes were recently discovered, involved in new defense systems with functions of RNA editing and satellite DNA retron synthesis, which are generically known by the surname of the first author of this work, Gao systems (27). Phages themselves are sometimes involved in the mobilization of these defense systems, and may even carry hotspots with several of them, which can make them interact positively and negatively with respect to other competing phages (17, 28).

Phages, in turn, defend against the bacterial defense systems with anti-defense systems in an ongoing arms race (29, 30). In addition, phages can co-opt bacterial antiviral systems, promoting their dissemination while using them to defend against other phages and increase their competitiveness (31).

Given that the different systems of the defensome of a bacterial species are found in its accessory genome, a representative number of genomes is required to perform the study at the pangenomic level. Bacteria usually maintain their accessory genes only if they provide an advantage in the environment in which they are found, otherwise they would act as useless ballast, especially in large populations that allow for rapid counterselection (32). Thus, if bacterial cells are subject to selection pressure by a phage from its environment, they will tend to maintain specialized defense systems against it. Moreover, if bacteria in turn has accessory genes that facilitate phage infection, such as membrane proteins that act as adhesion sites or entry receptors, the maintenance of these defense systems becomes even more important (33).

Here we have studied the defensome of the opportunistic pathogenic bacterium *Acinetobacter baumannii*, from a previous pangenome of almost 9,000 genomes. This bacterium has been designated a critical priority by the WHO, due to its high rate of antimicrobial resistance and the urgent need to find new drugs against it. We have found 81 different phage defense systems, the most common being the SspBCDE and Gao_Qat systems, which appear in the genomes most frequently isolated in hospital infections, global clone 2 (34), while the rest of the clonal groups replace the SspBCDE with R-M systems and have a greater number of integrated prophages in their genomes. Likewise, by means of machine learning we have found species-specific prophages that are avoided by certain defense systems, but also others that co-appear with certain defense systems. All this suggests that these systems would be used in fights between phages, which use bacteria as a battlefield.

## Materials and Methods

### Pangenome construction

The pangenome used was derived from 9,696 genomes belonging to the *A. baumannii* species in Rubio et al., 2023 (33). Each of these genomes has been structurally annotated, along with the reference genes inherent to the pangenome.

To eliminate redundancy, the skani tool v0.2.2 was used to calculate the Average Nucleotide Identity (ANI) by comparing all strains with each other. Subsequently, the groups of strains that shared 100% identity and coverage at the nucleotide sequence level were detected and eliminated, leaving only one strain as the reference of the group.

Of these remaining genomes, we discarded those that have a poor quality as *A. baumannii* genome. To do this, we used the number of genes shared with the rest of the *A. baumannii* strains, in addition to the phylogenetic distribution. All genomes that had a number of average shared genes lower than 2.650 in the pangenome and/or belonged to an outgroup branch of our phylogeny were removed from the pangenome. Thus, 8,929 genomes were kept.

### Defense system prediction

Defense systems were predicted in all genomes of *A. baumannii* using Defense Finder version 1.0.9 (35). Completed systems were considered, but isolated proteins containing HMM domains associated with defense systems (e-value ≤ 1e-5) were also included in the analysis. Once the antiviral systems of *A. baumannii* were obtained, the defensome was constructed by mapping the genes that compose each system with reference genes in the pangenome. Genes found in more than 99% of the strains were considered core genes present in virtually all genomes of the species and therefore would not produce differences across genomes.

For further analysis, we used only those defense systems present in at least 1% of total strains in some cases and the most frequent systems in others, to characterize the generalities from the bacterial species.

### Prophage prediction

Prophages were predicted in all genomes of *A. baumannii* by using Phigaro version 2.3.0 with default parameters and the *abs* mode (36). Only prophages with a sequence longer than 9 Kb were considered. All obtained prophages were clustered by similarity using MeShClust version 3.0 program with the identity threshold of 0.90 and the value of total initial sequences as the parameter –*v* and ¼ of total initial sequences as the parameter *–b*.

To combine similar prophage genomes with the sequence oriented in the two possible reading directions, we performed a similarity search with BLASTN (37) by comparing the reference sequences of each cluster with each other. Then, clusters whose reference sequence shared at least 90% identity and coverage in both sequences were combined. In addition, if a reference sequence from a cluster shared at least 95% identity and 100% coverage with larger reference sequence from another cluster, these clusters were also combined, to join possible fragments from the same prophage.

Using this protocol, we were able to identify a total of 2,144 prophages. To analyze all prophages and discard potential false positives, we considered prophages present in 0.1% of the strains (present in at least 9 genomes), leaving a total of 357 prophages. For further analysis, we used the prophages that were present in at least 10% of the strains of each major phylogenetic group (MLST) of interest. Thus, a total of 51 prophages were obtained, to which the phage DgiS1 was added (38).

To avoid redundancy, the prophage sequences were also subjected to an ANI analysis using skani, followed by a similarity check by BLAST (Suppl. Fig. S1). Thus, we combined the redundant prophages, resulting in a total of 351 different prophages (present in 0.1% of the strains), of which 47 were selected as the most frequent (present in at least 10% of the genomes of each major phylogenetic group).

### Molecular phylogeny

To construct the molecular phylogeny, the amino acid sequences of genes present in 99% of the analyzed genomes were selected. This resulted in a total of 589 core genes. The sequences were then joined by strain and aligned with the MAFFT v7.271 program using the E-INS-i option (39), which is suitable for alignments with a large number of strains and gaps. Subsequently, the alignment was refined by removing areas with numerous gaps, few informative sites, and duplicate sequences using the ClipKIT v2.2.4 algorithm with the KPI method (40). The molecular phylogeny was created using the IQ-TREE v2.3.4 tool with the VT+R10 model and a bootstrap of 1000 (41). The optimal model was calculated using the integrated ModelFinder algorithm in IQ-TREE. The phylogeny was represented using the R package ggtree v1.10.5 (42).

### Defense system-prophage associations

The aim of this study was searching for positive and negative associations between prophages and defense systems, so we wanted to have an accurate classification of defense systems according to what prophages were integrated in genomes.

Thus, predicted defense systems and prophages were used as input data in the built machine learning (ML) model. Only the most frequent defense systems were used as labels, while the 47 most prevalent prophages were split into training and testing set by a 7:3 ratio.

ML model was built and tested using R version 4.3.2. We computed XGBoost models using xgboost package version 1.7.7.1. We proposed a binary classification exercise in order to obtain importance, as a score that indicates how useful or valuable each feature was in constructing the improved decision trees within the model. A grid search was designed to determine the best parameters for eta (using 1e-3, 1e-2 and 1e-1), max_depth (1 to 10 with a step size of 5), min_child_weight (1 and 5), subsample (0.5 to 1 with a step size of 0.15), colsample_bytree (0.5 to 1 with a step size of 0.15), the hyperparameters for regularization gamma (from 1e-3 to 5 with increments of ten), lambda (0 and 1e-3 to 10, with increments of ten), and alpha (0 and 1e-3 to 10, with increments of ten), and using “binary:logistic” as objective. Model hyperparameters were tuned using validation data through cross-validation. When we increased the number of combinations of hyperparameters being tested, the model performance on the testing data set improved a little, since we obtained a better prediction score kappa (Suppl. Table S1).

### Model evaluation

Ten-fold cross validation was performed to assess the accuracy and prediction capacity of the model and avoid overfitting. The dataset was divided into 10 equal parts: one part was used for validation, and the remaining part for training. In the ten rounds, every part appears in the training and validation dataset, and model was trained and validated using all the 10-mers for validation. The bias of the generated model was observed by tracking the average accuracy over the ten-validation set.

### Protospacer search

The set of spacers were extracted with CRISPRCasFinder 4.2.20 (43) from the available genomes. These were compared against prophage sequences, for proto-spacer search, using the BLASTN 2.9.0+ algorithm, using the blast-short option for short sequences and a threshold of 90% percent spacer identity and coverage.

## Results

### *A. baumannii* genomes have 81 different defense systems

To study the defense systems against phages infecting *A. baumannii*, 8,929 bacterial genomes isolated worldwide were used. These genomes constitute a pangenome of 39,947 genes, with 2,220 core genes and 37,727 accessory genes.

Based on this pangenome, 81 different complete defense systems were found, with all but 98 genomes analyzed having at least one of them. These systems can be classified into 4 groups according to their mechanism of action: innate, adaptative, abortive or toxin/antitoxin, and unknown systems (Fig. 1A). The systems most present in the defensome of *A. baumannii* were the innate ones (present in 94% of genomes), followed by systems with unknown mechanisms (present in 51% of genomes)

**Fig. 1.**
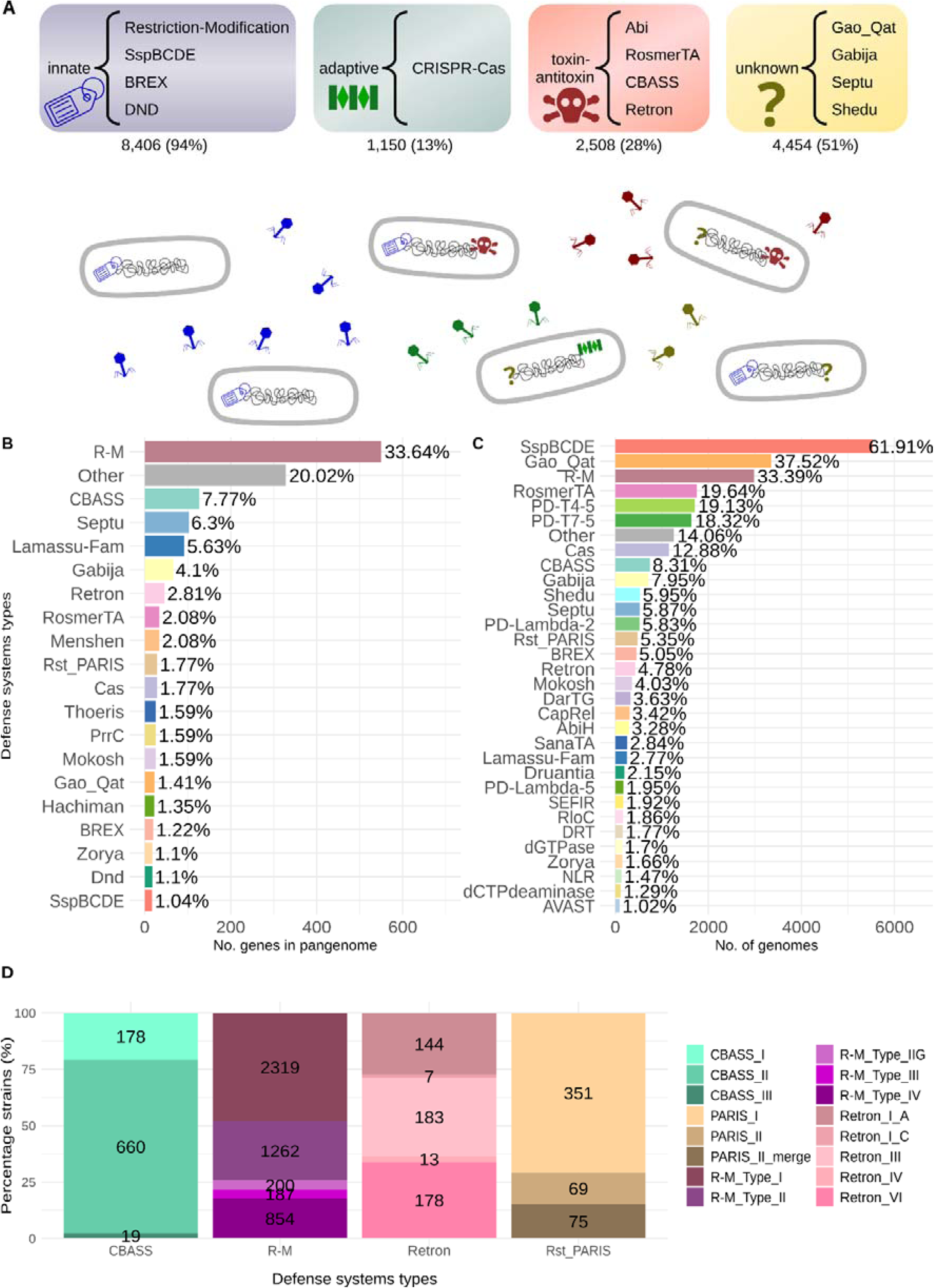
*A. baumannii* defensome. (A) Most frequently encountered defense systems grouped by type: innate, adaptive, toxin-antitoxin and unknown. Below each group is the number of genomes that have any system of this type, as well as the relative frequency of genomes. These results show that not all genomes have all systems, which are likely to be specialized in different phages. (B) Absolute and relative frequency of genes belonging to each system with respect to the total number of defense genes in the pangenome. (C) Absolute and relative frequency of genomes that possess each one of the systems, ordered from most to least frequent. (D) Number of genomes having each defense system subtype, for some of the systems.

In total, 1,619 different genes were involved in the different defense systems (4% of the pangenome). These are mainly accessory genes that do not appear in all strains and constitute the pan-immune system or defensome of the species (Suppl. Table S2). Only 11 core genes (*prmB*, *fokIM*, *cysH*, *yjjV*, *fstK*, *pkn5*, *htpG*, *mdcH 2*, *air*, *eptA 3*, *mgtA 3*), present in virtually all genomes, were predicted to assist certain very rare configurations of the defense systems BREX, CapRel, Cas, Mokosh, PD-Lambda-5, R-M, SspBCDE and Gao_Qat.

The R-M systems presented the largest collection of different genes, constituting 33.64% of the defensome (550 genes). However, only 33.39% of the genomes analyzed had a full version of either of the subtype of these innate systems (Fig. 1BC). In fact, none of the defense systems found appeared in all *A. baumannii* genomes. On average, each individual genome had an average of 3±1.9 different defense systems. The other innate system, SspBCDE, was composed of a total of 17 genes and appeared in 61.91% of the genomes analyzed, making it the most frequent of all defense systems. Other lesser-known defense systems, such as Gao_Qat, appeared in 37.52% of the genomes, suggesting its possible relevance in this species.

It should be noted that R-M systems, as well as the CBASS and Retron toxin-antitoxin systems, were subdivided into different types and subtypes, with for example 2,319 genomes with R-M systems type I and only 187 with type III, or 854 with type IV (Fig. 1D).

### Different innate defense systems do not usually appear together in the same genome

Since the different defense systems of *A. baumannii* are part of its accessory genome, we wanted to know if these were randomly distributed or if there was any coincidence or incompatibility of systems in the same genome.

By looking at pairs of systems that appear frequently in the same genome, the first thing that stands out is that most systems co-appear with R-M systems, resulting in the fact that when one of these systems appears, an R-M system also appears (Fig. 2A). Particularly noteworthy are the CRISPR-Cas and RosmerTA systems, which appear with high frequency when the genome also has an R-M system. The systems with minor occurrence when R-M systems appear in a genome are SspBCDE (the other innate system), Gao_Qat, Shedu and PD-T systems (PD-T4-5 and PD-T7-5, which appear together 3 out of 4 times).

**Fig. 2.**
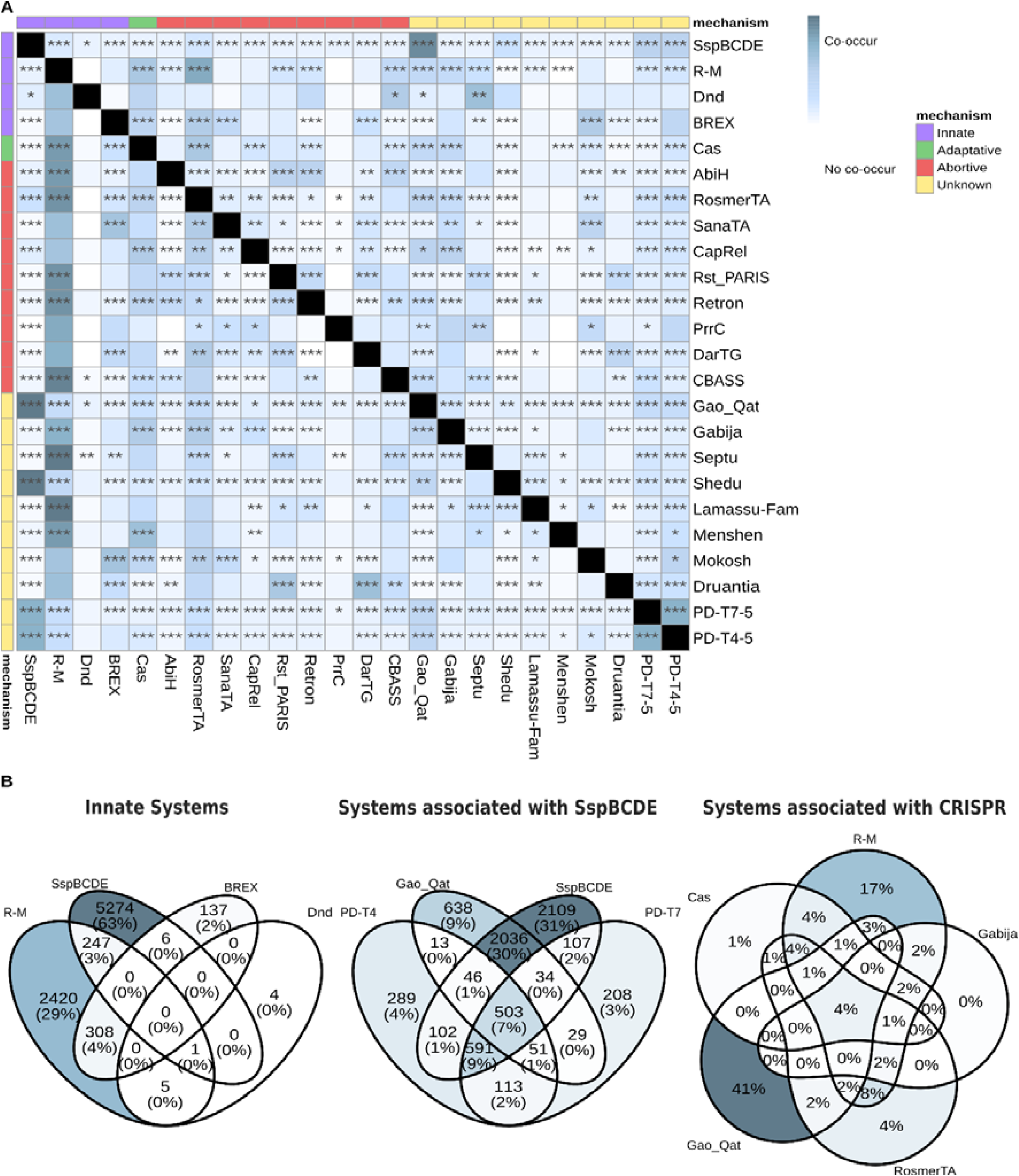
Co-appearance of defense systems. (A) Co-appearance of defense systems that appears in at least one percent of strains from the pangenome (97 genomes). The diagonal above shows the proportion of genomes that having the row system also has the column system, and the diagonal below shows the proportion of genomes that having the column system also has the row system. Fisher’s test was used to determine whether there is a significant association (positive or negative) between two systems (p-value < 0.001, ‘***’; p-value < 0.01, ‘**’; p-value < 0.05, ‘*’). (B) Multiple co-occurrences of sets of defense systems: innate systems, systems frequently associated with SspBCDE and systems frequently associated with CRISPR-Cas. In the first two cases the number of genomes is shown together with the relative frequency, and in the last case only the relative frequency is shown for simplicity.

Notably, the four innate systems of *A. baumannii* appear to be mutually exclusive, especially the R-M and SspBCDE systems (Fig. 2B), with the latter appearing strongly associated with the aforementioned systems not coappearing with the R-M systems: Gao-Qat and PD-T systems. However, the R-M systems present a high coappearance with the CRISPR-Cas systems, and 84% of genomes that have CRISPR-Cas systems also have R-M systems. In addition, genomes having CRISPR-Cas systems also tend to have RosmerTA (56%), Gabija (37%) and Gao-Qat (34%) systems with a high frequency (Fig. 2B). Since the R-M systems are known to synergize with CRISPR-Cas (11, 12), some of the other three mentioned systems could also help them in some of their steps.

### R-M systems appear to be more efficient than SspBCDE in avoiding phage integration

The defense systems found allow the bacteria to defend themselves against phage infection. To test their expected efficiency and specificity, we searched for phages integrated into the genomes of *A. baumannii*. Of the more than 2,000 different prophages found in the pangenome, only those that were present in at least 0.1% of the total number of genomes (group ‘total prophages’) and that appeared most frequently in the major clonal groups of the species (group ‘frequent prophages’) were taken for further analysis. These sets of prophages were searched in all bacterial genomes.

The first thing to note is that there was an average of 2.83±1.55 total prophages per genome (Fig. 3A). However, 183 genomes did not have any prophage and 1,613 did not have any of the 47 frequent phages. Most of these 183 genomes have R-M systems (60%), which suggests that these would be the most efficient systems against phages in the bacterium (Fig. 3B). Of the remaining genomes lacking prophages, 43 had the alternative innate SspBCDE system (with only 3 of them also having R-M systems), and 9 had innate BREX systems (with 5 genomes also having R-M systems). This would suggest that the R-M and SspBCDE systems would be alternative innate systems, while the BREX systems may be synergistic to the R-M systems. Remarkably, 7 of the genomes without prophages do not exhibit any defense system. This suggests that they either have as yet undescribed defense systems or are bacteria that have not been under selection pressure from phages.

**Fig. 3.**
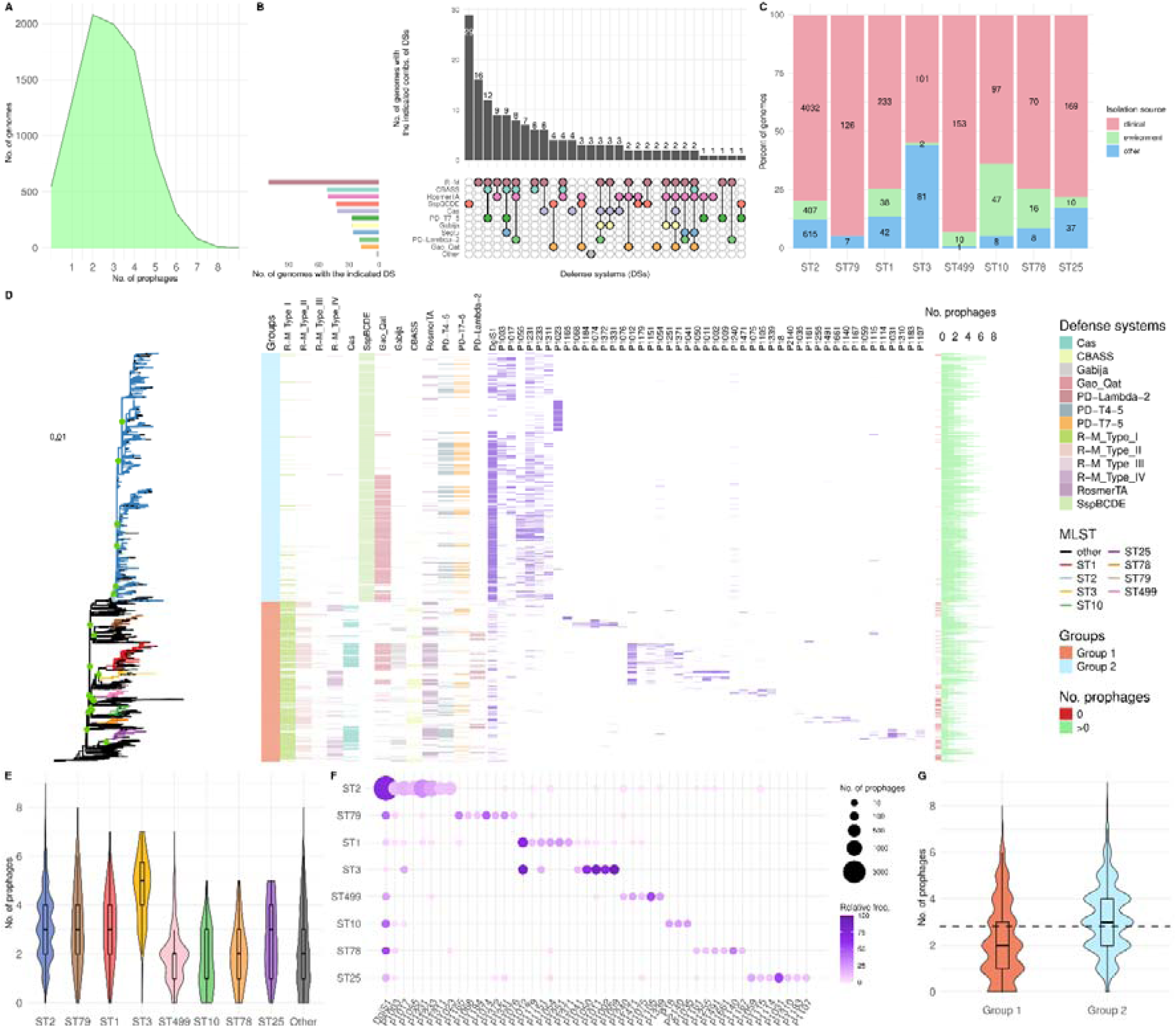
Prophages in the genomes of *A. baumannii*. (A) Number of genomes that have a given number of total prophages. (B) Number of prophage-lacking genomes differentiated by the defense systems they carry and their combinations. (C) Number of genomes of each of the most frequent clonal groups of the species, differentiating by their isolation site. (D) Molecular phylogeny of all the genomes analyzed, highlighting in the tree the most frequent clonal groups. Nodes separating the main MLST groups with a boostrap value higher than 80 are shown with green dots. From left to the right, it shows in order: distribution of the two main groups in the phylogeny (group 1 and group 2 genomes), presence-absence of the most frequent defense systems, presence-absence of the most frequent prophages by clonal group and number of total prophages (0 prophages is shown in red). (E) Distribution of the number of total prophages in the different clonal groups. Significant differences between the numbers of phages in each clonal group were determined using the Kruskal-Wallis test (Kruskal-Wallis, p-value = 4.01e-214) (F) Presence-absence (abundance) of the most frequent prophages in the different clonal groups. (G) Distribution of the number of total prophages in the two main groups of genomes (group 1 and group 2) shown in the phylogeny with the average number of total prophages. Significant differences between the numbers of phages in each clonal group were determined using the Wilcoxon test (Wilcoxon, p-value = 2.63e-141).

The defense systems of a bacterial species are part of what is known as the pan-immune system. In support of this idea, the same system should appear scattered throughout the pangenome of the species, and specific defense systems could be gained or lost by genomes of the species, with the global population being a reservoir for all of them. However, certain defense systems could be linked to clonal groups of the bacterium, if these offer them some advantage in their environment, such as defense against the phages they may encounter there. To analyze this, we studied the distribution of the most frequent defense systems according to the phylogenetic relationships of the genomes containing them, with special emphasis on the most frequently isolated clonal groups of *A. baumannii*, which have been mainly isolated in clinical environments (Fig. 3CD). In addition, we analyzed the presence/absence and abundance of frequent prophages according to their phylogenetic relationships and their association with frequent defense systems, to test the effectiveness of defense systems.

The prevalent ST2 clonal group and other phylogenetically close groups (group 2) seem to have almost exclusively the SspBCDE system in their genomes (96%), while the rest of the groups, evolutionarily distant from ST2, present R-M systems instead (group 1). Group 2 also presents, with less frequency, the other defense systems seen as associated with SspBCDE, the most common being the Gao-Qat and the PD-T systems (45% and 26%, respectively). Meanwhile, the rest of the genomes present a greater heterogeneity of defense systems, although always lacking the SspBCDE system. This correlates with the fact that there are more prophages on average in ST2 (group 2) than in other less prevalent clonal groups (Fig. 3E). One of the most frequent prophages in this clonal group is DgiS1 (Fig. 3F), which we previously proved competes with a bacterial CRISPR-Cas I-F system for the same integration site (38). If we consider that the defense island in which the prophage is integrated is also the site of integration of the genes of the Gao_Qat system, this could point to the fact that these defense systems could be associated with specific prophages of the bacterium, which would use them as weapons to fight against other different prophages. In fact, genomes that present an SspBCDE or Gao_Qat system as the only defense system presented 3-4 integrated prophages (more than the average of all genomes). Another prophage in the ST2 group is P1023, which appears in genomes with SspBCDE, but not with Gao-Qat or PD-T.

On the other hand, although many of the different prophages of the species appear in different clonal groups, there are prophages that have a greater frequency in certain taxonomic groups, and even almost exclusively in some of them. This shows how clonal groups with specific combinations of defense systems tend to have different prophage profiles. For example, prophage P1012 in ST1 group, prophage P1031 in the ST25 group, prophages P1009 and P1011 in ST3 group (plus prophage P1012), and prophage P1105 in ST499 group (Fig. 3F).

Genomes in group 1 do not appear to have a high proportion of CRISPR-Cas and CBASS systems (1045 and 612 genomes, respectively), co-occurring only in 148 genomes. Of those with CRISPR-Cas, 83% also had R-M systems, and of those with CBASS, 86%. However, the alternative innate SspBCDE system did not co-appear with CRISPR-Cas systems in any of these genomes, and only in 5 genomes did it co-appear with CBASS systems across the entire pangenome.

These results suggest a highly differentiated defense system profile between the ST2 and similar clonal groups (group 2) and the rest of the *A. baumannii* clonal groups (group 1), with the SspBCDE system being specific to group 2 and the R-M systems to group 1.

These two groups show distinct prophage profiles and also show a quantitative difference in the number of prophages, with an average of 2.18 ±1.63 prophages in group 1 and 3.12±1.43 prophages in group 2 (Fig. 3G). This could be due to the homogeneous profile of the defense systems or to the low number of systems in this clonal group 2.

### The presence of certain defense systems is associated with specific prophages

To evaluate the efficiency of the defense systems to prevent phage integration into the bacterium, the number of total prophages present in genomes with a given system was tested versus those without. Interestingly, the most abundant systems (SspBCDE and Gao-Qat) appeared with a higher number of prophages (average higher than 3), while the 98 genomes lacking any defense system had an average of 2 prophages (Fig. 4A).

**Fig. 4.**
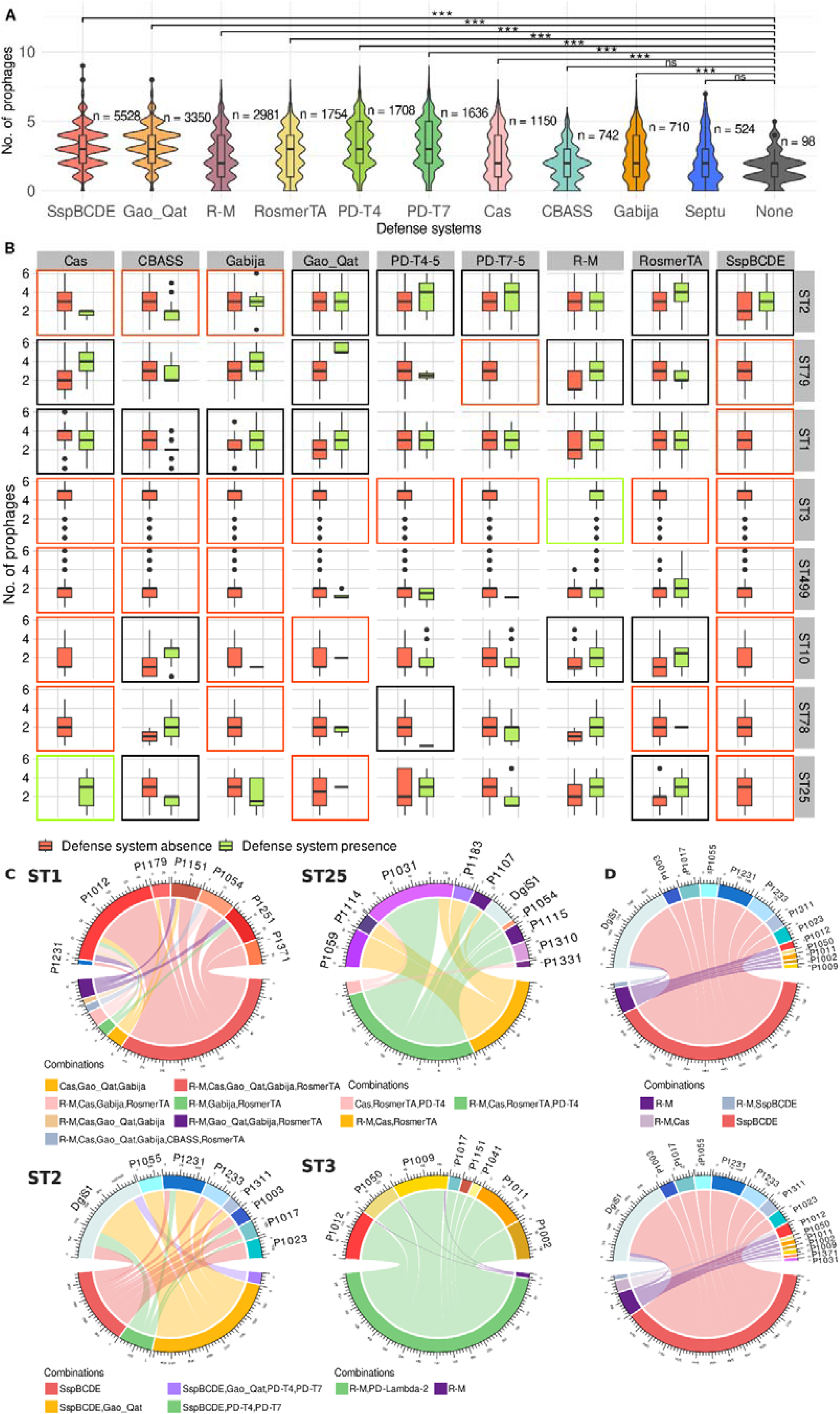
Number of prophages according to the defense system possessed by the bacterium. (A) Distributions of the number of prophages according to the defense system of the bacterium (Paired Wilcoxon-test with the group ‘None’; p-value < 0,001, ***; ns, non-significant). (B) Distributions of the number of prophages in the bacterium separated by clonal groups and differentiating whether they have or do not have certain defense systems. A black box indicates that there is a significant difference, red if there are only genomes with absence of the defense system (or there are less than 1% of genomes per each clonal group with presence), green if there are only genomes with presence (or there are less than 1% of genomes per each clonal group with absence), and the absence of a box indicates that there is no significant difference (Wilcoxon-test, p-value < 0.05). (C) Specific prophages occurring in bacterial genomes with specific combinations of defense systems. Each individual circos is constructed from a single ST clonal group, indicated in the figure, using only those combinations found in at least 2% of the genomes of such clonal group (this threshold was used for clarity). The numbers around the circos indicate the number of genomes found with that prophage and that combination of defense systems. (D) The same than (C), comparing only the innate defense systems SspBCDE and R-M throughout the entire pangenome (including all ST groups), using only those combinations found in at least 1% of all genomes. Above: genomes with R-M or SspBCDE as the only defense system; Below: genomes with R-M, R-M and CRISPR-Cas, or SspBCDE as the only defense system.

However, all these calculations could be biased by the different representation of *A. baumannii* clonal groups. Therefore, it was decided to perform this evaluation by separating again the genomes by the most frequent clonal groups. Thus, if we consider that a clonal group is a group of genomes that are taxonomically very close, small changes in them, such as the presence of a certain defense system could be associated with a lower number of prophages, if that system is efficient against certain phages.

For the CRISPR-Cas systems, it was observed that in the ST25 group there were only genomes with this defense system, while in the ST10, ST2, ST3 and ST499 groups this system does not appear (Fig. 4B). However, in the ST79 clonal group there are genomes with and without CRISPR-Cas system, with a significant difference in the number of prophages present, in favor of the presence of the defense system. Specifically, CRISPR-Cas systems are present in genomes with an average of 3 prophages, while the absence of the defense system occurs in genomes with an average of 2 prophages. The presence of a higher number of prophages also occurs in ST2 group when PD-T, RosmerTA and SspBCDE systems are present.

In the ST1 clonal group, CRISPR-Cas system presence is related with a lower number of prophages, indicating the effectiveness and explaining the prevalence of this system in this clonal group. A fewer number of prophages is also associated with the presence of CBASS and RosmerTA in particular clonal groups.

Focusing on the innate systems, SspBCDE systems are associated with a higher number of prophages, as mentioned in ST2 group. And, in clonal groups where R-M systems predominate, there is no significant difference between the number of prophages when R-M is present or absent, except in the case of ST79 and ST10 groups where again the presence of R-M is associated with a higher number of prophages in the bacterial genome.

Differences in the number of prophages depending on the presence or absence of a certain defense system could be due to the presence or absence of specific prophages (sensitive or resistant to that defense system). To test this, the presence of the most frequent prophages in each clonal group was evaluated versus the combination of defense systems in their genomes (Fig. 4C). Thus, in the ST1 group, the CRISPR-Cas system is associated with the presence of many of the most frequent phages in this clonal group. Therefore, the quantitative difference between presence and absence of the CRISPR-Cas system that can be seen in Fig. 4B would be due to the rest of prophages that are not frequent in this group.

Another case is the ST25 group, in which the presence of the PD-T4 system is associated with the presence of 7 different prophages, and at the same time is clearly associated with the absence of prophages P1059, P1114 and P1183. In the ST2 group, each different combination of defense systems is related to the presence of different phages, except in the case of the DgiS1 prophage, which always appears, regardless of the defense systems of the bacterium. And in the ST3 group, the absence of the PD-Lambda system is related to a lower occurrence of the most frequent prophages.

On the other hand, in the ST79 group, the presence of CRISPR-Cas systems is associated with the presence of 6 different prophages, whereas prophage P1165 does not appear together with the defense system (Suppl. Fig. S2). This suggests that the CRISPR-Cas system may not prevent the presence of certain prophages, but it may prevent the presence of the aforementioned prophage P1165, which is not avoided by the R-M systems.

Other interesting cases are the ST10 and ST78 groups. In the ST10 group, the presence of PD-T systems seems to prevent the occurrence of most of the frequent prophages in that group. This fact is reflected in the differences in the total number of prophages in genomes with the presence of the aforementioned systems in this clonal group, with a lower number of prophages being observed when the bacteria are armed with these systems (Figure 4B). Finally, in the ST78 group, the presence of the PD-T and Gao_Qat systems seems to prevent the infection of most of the common prophages, as can be seen in the total number of prophages.

Furthermore, when comparing the collection of prophages present and absent in genomes having only one defense system of the two innate R-M (with or without CRISPR-Cas) and SspBCDE, the prophage profile they provide is completely different (Fig. 4D).

### Prophage combinations allow prediction of which defense systems a bacterium carries

Finally, having seen that certain prophages appeared negatively or positively related to certain defense systems (or their combinations), we wanted to know if the presence-absence pattern of prophages could be used to predict if a genome had a certain defense system. To do this, we created a predictive model, using machine learning techniques, to calculate the importance or weight of each individual prophage in tagging a bacterial genome with a given defense system. This importance could be due to two possible relationships: negative (the prophage is rare when the defense system appears, and therefore suggests that the system avoids prophage infection), or positive (the prophage is very frequent when the defense system appears, and therefore suggests that the system does not avoid phage infection, or the phage could be the carrier of the defense system).

The results of this experiment showed that few prophages can be related to a given defense system, i.e., most prophages could be avoided by any defense system under certain circumstances (Fig. 5A). In the case of CRISPR-Cas systems, the low frequency of three prophages (P1011, P1017 and P1023) combined with the presence of two others (P1012 and P1031) allows us to predict with high confidence that this defense system is present. The presence of prophage P1012 is also important in predicting that the bacterium is armed, in addition to CRISPR-Cas, with the R-M, Gabija and RosmerTA systems, which are three defense systems that co-occur with high frequency with CRISPR-Cas, as shown above. In addition, the lack of prophage P1023 is also important in predicting that a genome has R-M systems, as well as PD-T systems, RomerTA and CBASS In contrast, the presence of the prophage is important for predicting that the bacterial genome has the other innate system, SspBCDE.

**Fig. 5.**
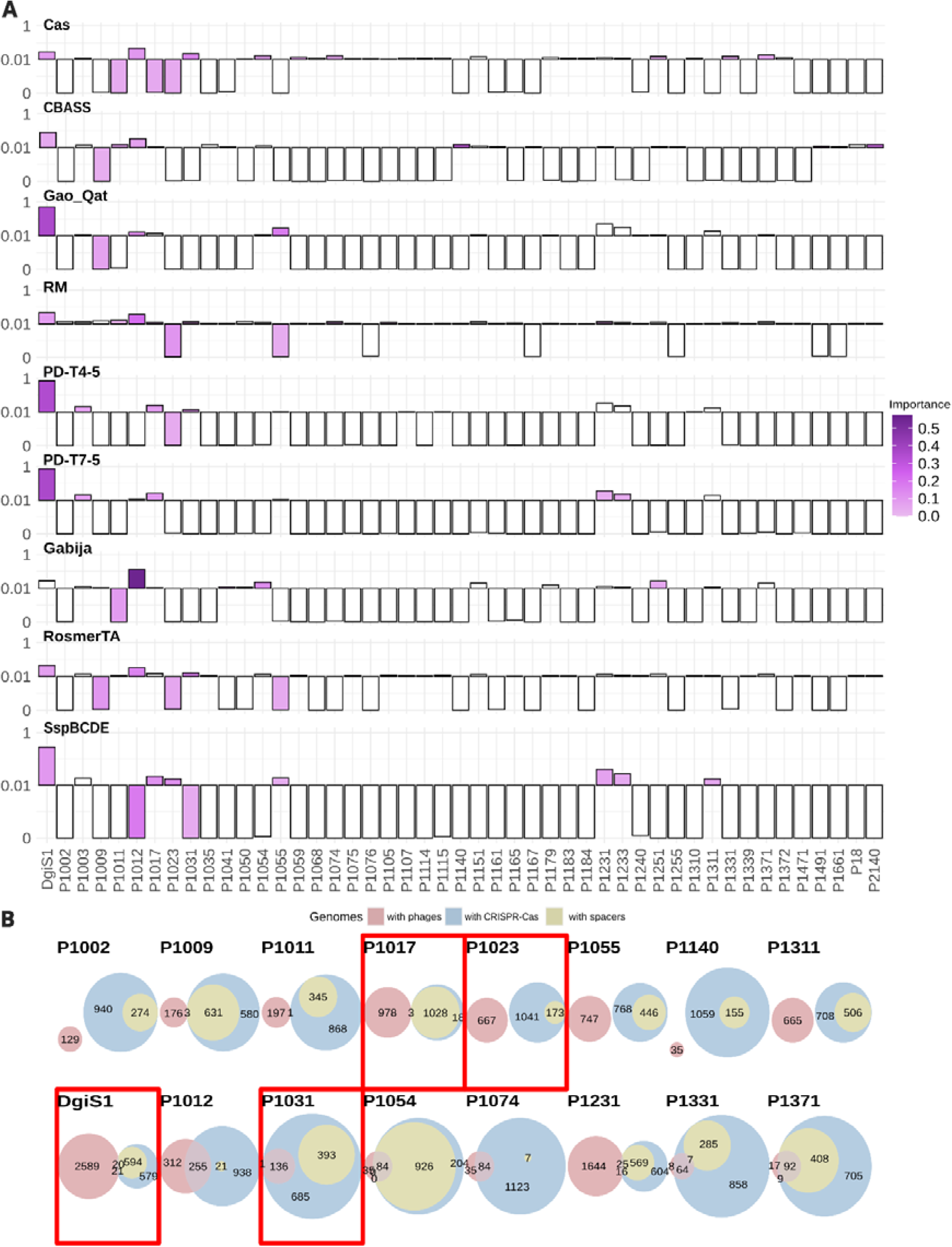
Machine learning approach to search for important prophages related to a particular defense system. A) Importance of the different prophages in the classification of a genome as having a particular defense system. For each defense system, the bar color shows the importance of each prophage (from 0 to 0.57, as maximum importance in all defense systems) in the classification of a genome as having that system. Relative abundance shows whether a given prophage is very abundant (bars up) or low abundant (bars down) in genomes that have a given defense system (if the relative abundance is lower than 0.01, this abundance is multiplied by 5 and subtracted by the maximum abundance of each type, to get negative values and increase the differences between them). B) Number of genomes that have a given prophage, highlighting those that have only the prophage, only CRISPR-Cas or CRISPR-Cas systems with spacers against the prophage, and combinations of all of these. The top row presents all prophages negatively correlated with the CRISPR-Cas system (importance > 0.005 and relative frequency <= -0.98) and the bottom row those positively correlated (importance > 0.01 and relative frequency >= 0.05). The Venn diagrams highlighted by a red box represent the prophages shown as important by machine learning.

In fact, the two main innate systems present an almost inverse prophage profile, as previously seen. Specifically, the presence of prophage DgiS1, as well as prophages P1017 and P1023, together with the absence of prophage P1012 are clearly associated with the presence of the SspBCDE system. However, the opposite trend in prophages P1012 and P1023 anticipates that the bacterial genome has instead an R-M system.

The prophage DgiS1 is a special case. It is a very frequent prophage in *A. baumannii*, which is not only important for predicting SspBCDE in the genome, but it also appears to be specially associated with the PD-T and Gao_Qat systems. PD-T systems are found inside the sequence of the prophage itself, while the second had already been found with a high frequency of co-occurrence previously, since this system shares integration site with DgiS1 (38). In the case of SspBCDE, the DgiS1 prophage does not carry the system itself, but it does bear the *sspD* gene of the defense system, which encodes an enzyme whose function is to add phosphorothioate groups, which could reflect a refined strategy to mark the phage genome and make it undetectable by the bacterial SspBCDE system. This gene was found in 63.6% of Dgis1 prophage genomes (expect value = 1.3e-14 with HMM profile).

The meaning of the results obtained by machine learning can be further probed in the case of a defense system, the CRISPR-Cas. In this case we can draw some predictions. If the absent phages are missing because they are fought away by the defense system, their CRISPR arrays must contain spacers that target the sequence of these prophages. Likewise, if the prophages are associated with the presence of the defense system, we should not find spacers, or if found, they would not prevent the presence of the prophage.

To verify this fact, we checked the number of genomes in which specific prophages appeared, together with the CRISPR-Cas systems and spacers that target the prophage in those systems, as well as the overlap of all of them. The result obtained was that in the case of prophages negatively associated with CRISPR-Cas systems, the genomes in which the prophage appears have no CRISPR-Cas systems, and when this system appears the prophage is not present but the CRISPR arrays usually present spacers against the prophage (Fig. 5B). However, in the case of prophages positively associated with CRISPR-Cas systems, the prophage appears to coexist with this defense system, and either the CRISPR arrays do not contain spacers facing the phage, or if they do (as in the case of prophages P1031, P1054, P1331, and P1371), the prophage and spacers usually do not coincide in the same strain. This is so, except in the case of prophages P1054 and P1371, where in many cases the prophage and the spacers against it co-appear in the same bacterial genome. Of these last two prophages, the prophage P1371, precisely, presented a relative high importance for classifying a genome as having CRISPR-Cas systems.

## Discussion

It is estimated that phage infections cause a daily bacterial mortality rate of 5-50% (2, 44). In turn, bacteria defend themselves by means of multiple antiphage defense elements that are distributed heterogeneously throughout their genomes (45, 46). Specifically, here we have found 81 antiphage defense systems in a pangenome of the species *A. baumannii*, none of them being found in all of the nearly 9,000 genomes analyzed. However, these systems are probably available in the species, as a pan-immune system, being maintained in specific strains of environments where they suppose a competitive advantage (18). This idea is supported by the fact that we have sometimes found a different prophage profile among different clonal groups of the bacterium, associated with a certain profile of defense systems. Among the defense groups found, almost two thirds of the genomes have the innate SspBCDE system, sometimes coincident and sometimes not with Gao-Qat and PD-T4 systems, which are more frequent in this group. Although this high number could be partly influenced by the high frequency of isolation and sequencing of members of the ST2 clonal group, which has this combination of defense systems, from hospital infections. Meanwhile, the remaining genomes replace the SspBCDE system with R-M systems, combined with a heterogeneous number of other defense systems. All this would support the idea that systems are continuously gained and lost by bacterial genomes (47).

Specifically, the genomes of *A. baumannii* analyzed showed an average of 3±2 different defense systems, different from previous studies that using the same methodology found an average of 5 (35, 48). This difference could again be due to the high number of genomes from the abundantly sequenced ST2 clonal group, which are precisely those that possess a lower number of defense systems. In any case, the high number of unequally distributed defense systems throughout the *A. baumannii* pangenome would be correlated with the flexibility of this species (49), which could thus adapt to different environments dominated by different mobile genetic elements. Thus, some of these antiviral systems have a high frequency of coappearance, as we have found in the case of the *A. baumannii* CRISPR-Cas I-F and R-M type II systems. This fact has been described before and could be explained by a synergy that would improve the success of the former in creating new immunity to previously unrecognized phages (11, 12). Spacer adaptation in CRISPR-Cas systems can be enhanced in cells also carrying R-M systems, as these would limit phage replication through degradation of their genome, and these excision products can be captured by the adaptation machinery of these systems (11). Likewise, the coexistence of CRISPR-Cas and R-M systems leads to a decrease in the frequency of spontaneous phage mutants that escape these defense systems (50). All this makes CRISPR-Cas systems one of the most efficient defense systems in preventing the gain of mobile genetic elements in bacteria, including prophages (47), and this is particularly remarkable in the genus *Acinetobacter* (16).

The appearance of certain systems may be due to the environment in which the bacterium is found, with the presence of certain phages acting as a selection pressure. Similarly, the coappearance of systems may be due to the same environmental constraints, in addition to the functional synergy of certain aforementioned combinations. SspBCDE was the most frequently found defense system. This system appears positively associated with the prophage DgiS1 of *A. baumannii* and PD-T systems, precisely because the genes that conform the PD-T systems appear within the sequence of the prophage, which may be responsible for its dissemination in the species (38). The SspBCDE and Gao-Qat systems (the second most prevalent in the pangenome analyzed) are additional systems that appear to be positively connected to DgiS1. The Gao-Qat system is a defensive system that is found in the same defense archipelago where DgiS1 is integrated, next to a bacterial tmRNA gene. This may suggest synergy or symbiosis between both systems.

Thus, the innate SspBCDE and R-M systems in *A. baumannii* appear to have an antagonistic relationship. It is known that certain defense systems can show antagonistic relationships that prevent their stable coexistence, but the incompatibility can be overcome by epigenetic silencing, in which, precisely, these systems may be involved (51). Moreover, in this case, there would be two systems that label the bacterial DNA, so that only one of them would be sufficient for the proper functioning of an innate system. Here we have shown that the presence of these two defense systems is strongly associated with certain clonal groups of the bacterium, which also exhibit specific prophage profiles. In fact, the clonal group associated with the SspBCDE system has a higher number of prophages than the groups with the R-M system, suggesting that this ST2 clonal group, which is the most prevalent in hospital environment, could benefit from a more relaxed phage tolerance. This would lead to further infection and integration of new prophages, as these viral agents could contribute virulence factors or other defense systems to enhance the fitness and pathogenicity of the bacterium. It is interesting to note that the defense mechanisms possessed by these groups may actually contribute to their genetic divergence. For example, R-M systems have been shown to alter the rate of evolution in bacteria and to allow extensive genomic rearrangements and contribute to the genetic isolation of clonal groups (52, 53).

The R-M systems per se, considering all the types that encompass them, seem to be the innate system with the highest efficiency in avoiding phage infection, due to its prevalence in prokaryotes (5). Here we have found 550 different pangenome genes involved in R-M systems. This high number of genes suggests that it is a system exposed to constant evolution, making new variants appear that overcome the defenses of the phages themselves (54). The appearance of R-M systems, even without the coappearance of any other, causes the bacteria to have none or very low numbers of prophages.

In many cases it was seen that the presence of specific defense systems is associated with the presence of a higher number of prophages, which could be due in many cases to the fact that the phage can carry the defense system. This has been demonstrated in the nearby *Pseudomonas aeruginosa* species with some defense systems such as R-M type IV and Wadjet (47), and Abi and CBASS in other studies (16). Although in other cases it may be due to the fact that the phage has acquired immunity to the defense system, either because of some particular characteristic of the phage, or because it carries anti-defense systems (55). All this has been contrasted in the analysis performed by machine learning, being especially remarkable the fact that several prophages seem to be avoided by R-M and CRISPR-Cas systems, while others seem to positively correlate with their appearance. In contrast, the same prophages have the opposite tendency in genomes presenting SspBCDE. These results could be supported in the case of CRISPR-Cas systems, thanks to the analysis of the spacers. All this demonstrates the usefulness of this type of analysis in microbiology (56).

It is noteworthy that the genomes in which no defense system was found, the number of prophages was lower and close to 0. This could be due to the fact that these cells do not usually encounter phages in their environment, and therefore carrying defense systems in their genomes would be a burden. But it could also be associated with the fact that phages mobilize defense systems, and thus both elements correlate positively. However, this hypothesis will need to be demonstrated in the future, in large-scale evolutionary studies.

We have shown, by means of machine learning modeling, that by taking into account the prophage profile, it is possible to predict which defense system a given strain of bacteria will have. This information is relevant for the use of phages to treat bacterial infections or phage therapy (57). When analyzed for lytic phages, the antiviral systems in the oyster-colonizing bacterium *Vibrio crassostreae* have been shown to be important in determining the phage-host range (58). However, in *E. coli*, genes encoding membrane proteins that are used by phages as adsorption factors or for the entry into the bacterial cell, have been found to be more crucial in determining the bacterial strain-phage interaction, while the defensome would be less relevant in predicting this interaction (59). Using a different approach, we have shown here that antiviral systems could, indeed, modulate the bacterial phageome, considering the phages already integrated into the bacterial genome. All this, added to the defense systems that phages occasionally carry, could be shaping and constraining the mobilome of the species.

## Acknowledgements

We thank C3UPO for the HPC support.

## Data and Materials Availability

All data needed to evaluate the conclusions in the paper are present in the paper and/or the Supplementary Materials. The specific code and input data used are available at Zenodo (https://zenodo.org/records/14218265).

## Supplementary Files

**Suppl. Table S1.** Best hyperparameters and scores from the ML models (one for each defense system).

**Suppl. Table S2.** Defense systems and their genes found in *A. baumannii* pangenome.

**Suppl. Fig. S1.**
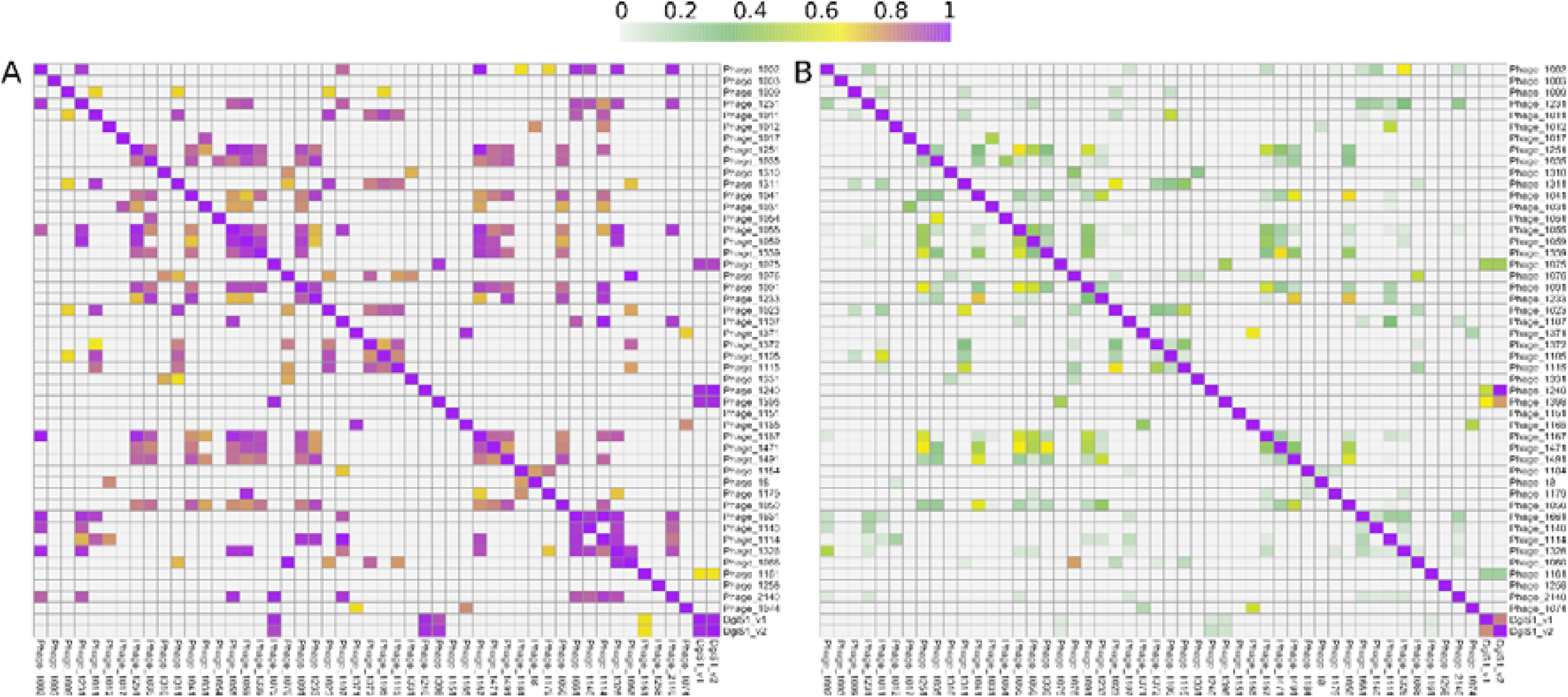
Average nucleotide identity (ANI) of the prophages found in the bacterial genomes.

**Suppl. Fig. S2.**
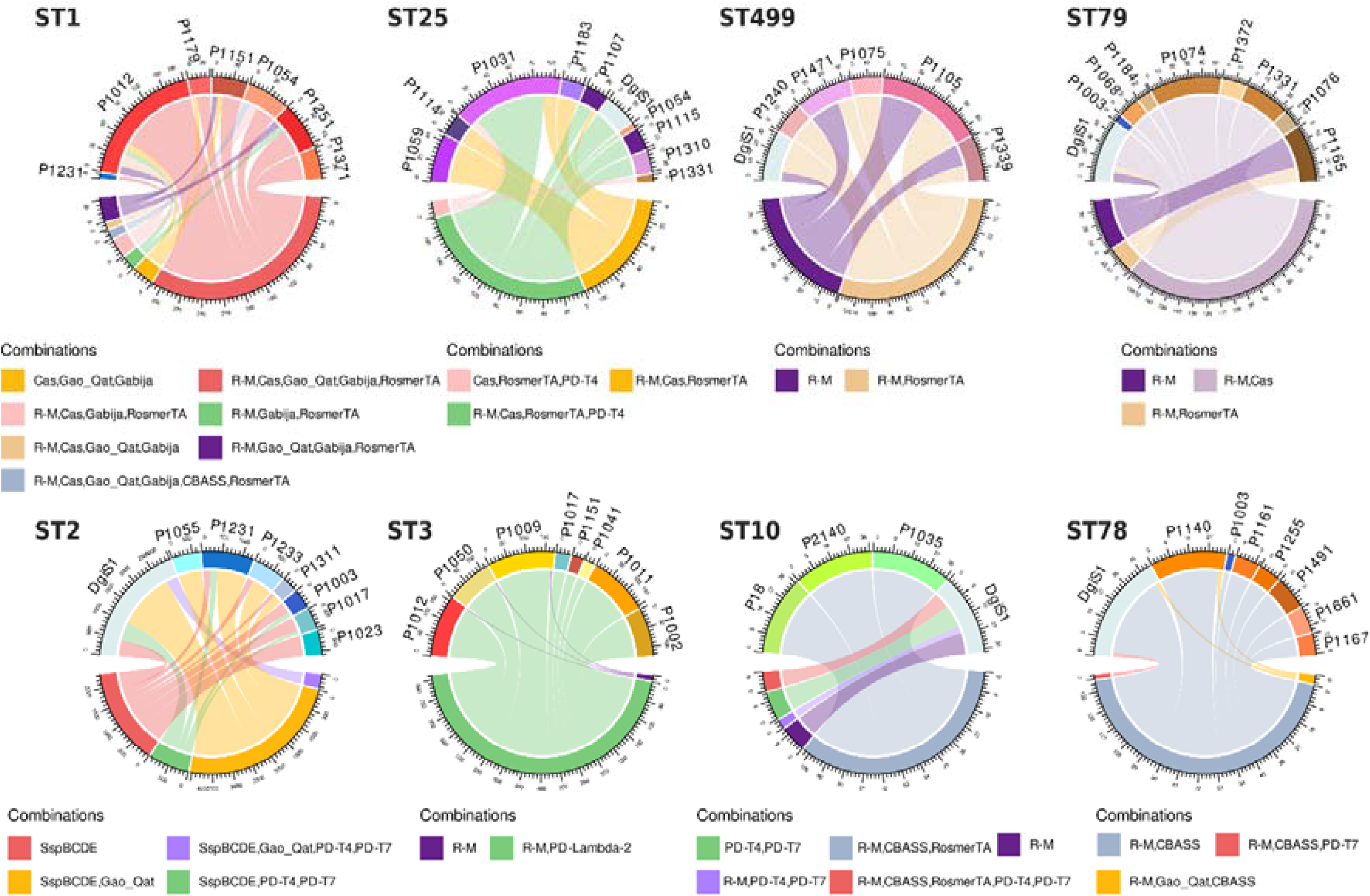
Specific prophages occurring in bacterial genomes with specific combinations of defense systems (clonal groups missing in Fig. 4C).

## References

1. Chibani-Chennoufi S, Bruttin A, Dillmann M-L, Brüssow H. 2004. Phage-Host Interaction: an Ecological Perspective. Journal of Bacteriology 186:3677–3686.

2. Fuhrman JA, Noble RT. 1995. Viruses and protists cause similar bacterial mortality in coastal seawater. Limnology and Oceanography 40:1236–1242.

3. Bertozzi Silva J, Storms Z, Sauvageau D. 2016. Host receptors for bacteriophage adsorption. FEMS Microbiol Lett 363:fnw002.

4. Labrie SJ, Samson JE, Moineau S. 2010. Bacteriophage resistance mechanisms. Nat Rev Microbiol 8:317–327.

5. Oliveira PH, Touchon M, Rocha EPC. 2014. The interplay of restriction-modification systems with mobile genetic elements and their prokaryotic hosts. Nucleic Acids Research 42:10618– 10631.

6. Roberts RJ, Vincze T, Posfai J, Macelis D. 2023. REBASE: a database for DNA restriction and modification: enzymes, genes and genomes. Nucleic Acids Res 51:D629–D630.

7. Nathans D, Smith HO. 1975. Restriction endonucleases in the analysis and restructuring of dna molecules. Annu Rev Biochem 44:273–293.

8. Mojica FJM, Díez-Villaseñor C, García-Martínez J, Soria E. 2005. Intervening sequences of regularly spaced prokaryotic repeats derive from foreign genetic elements. J Mol Evol 60:174– 182.

9. Bolotin A, Quinquis B, Sorokin A, Ehrlich SD. 2005. Clustered regularly interspaced short palindrome repeats (CRISPRs) have spacers of extrachromosomal origin. Microbiology (Reading) 151:2551–2561.

10. Sorek R, Kunin V, Hugenholtz P. 2008. CRISPR--a widespread system that provides acquired resistance against phages in bacteria and archaea. Nat Rev Microbiol 6:181–186.

11. Hynes AP, Villion M, Moineau S. 2014. Adaptation in bacterial CRISPR-Cas immunity can be driven by defective phages. Nat Commun 5:4399.

12. Dupuis M-È, Villion M, Magadán AH, Moineau S. 2013. CRISPR-Cas and restriction–modification systems are compatible and increase phage resistance. Nat Commun 4:2087.

13. Aframian N, Eldar A. 2023. Abortive infection antiphage defense systems: separating mechanism and phenotype. Trends Microbiol 31:1003–1012.

14. Duncan-Lowey B, Kranzusch PJ. 2022. CBASS phage defense and evolution of antiviral nucleotide signaling. Current Opinion in Immunology 74:156–163.

15. Kelly A, Arrowsmith TJ, Went SC, Blower TR. 2023. Toxin-antitoxin systems as mediators of phage defence and the implications for abortive infection. Curr Opin Microbiol 73:102293.

16. Liu Y, Botelho J, Iranzo J. 2024. Timescales and genetic linkage explain the variable impact of defense systems on horizontal gene transfer. bioRxiv 10.1101/2024.02.29.582795.

17. Fillol-Salom A, Rostøl JT, Ojiogu AD, Chen J, Douce G, Humphrey S, Penadés JR. 2022. Bacteriophages benefit from mobilizing pathogenicity islands encoding immune systems against competitors. Cell 185:3248–3262.e20.

18. Bernheim A, Sorek R. 2020. The pan-immune system of bacteria: antiviral defence as a community resource. Nat Rev Microbiol 18:113–119.

19. Millman A, Melamed S, Leavitt A, Doron S, Bernheim A, Hör J, Garb J, Bechon N, Brandis A, Lopatina A, Ofir G, Hochhauser D, Stokar-Avihail A, Tal N, Sharir S, Voichek M, Erez Z, Ferrer JLM, Dar D, Kacen A, Amitai G, Sorek R. 2022. An expanded arsenal of immune systems that protect bacteria from phages. Cell Host Microbe 30:1556–1569.e5.

20. Jiang S, Chen K, Wang Y, Zhang Y, Tang Y, Huang W, Xiong X, Chen S, Chen C, Wang L. A DNA phosphorothioation-based Dnd defense system provides resistance against various phages and is compatible with the Ssp defense system. mBio 14:e00933–23.

21. Goldfarb T, Sberro H, Weinstock E, Cohen O, Doron S, Charpak-Amikam Y, Afik S, Ofir G, Sorek R. 2015. BREX is a novel phage resistance system widespread in microbial genomes. EMBO J 34:169–183.

22. Wang S, Wan M, Huang R, Zhang Y, Xie Y, Wei Y, Ahmad M, Wu D, Hong Y, Deng Z, Chen S, Li Z, Wang L. 2021. SspABCD-SspFGH Constitutes a New Type of DNA Phosphorothioate-Based Bacterial Defense System. mBio 12:e00613–21.

23. Xiong X, Wu G, Wei Y, Liu L, Zhang Y, Su R, Jiang X, Li M, Gao H, Tian X, Zhang Y, Hu L, Chen S, Tang Y, Jiang S, Huang R, Li Z, Wang Y, Deng Z, Wang J, Dedon PC, Chen S, Wang L. 2020. SspABCD-SspE is a phosphorothioation-sensing bacterial defence system with broad anti-phage activities. Nat Microbiol 5:917–928.

24. Zou X, Xiao X, Mo Z, Ge Y, Jiang X, Huang R, Li M, Deng Z, Chen S, Wang L, Lee SY. 2022. Systematic strategies for developing phage resistant Escherichia coli strains. 1. Nat Commun 13:4491.

25. Hochhauser D, Millman A, Sorek R. 2023. The defense island repertoire of the Escherichia coli pan-genome. PLOS Genetics 19:e1010694.

26. Makarova KS, Wolf YI, Snir S, Koonin EV. 2011. Defense islands in bacterial and archaeal genomes and prediction of novel defense systems. J Bacteriol 193:6039–6056.

27. Gao L, Altae-Tran H, Böhning F, Makarova KS, Segel M, Schmid-Burgk JL, Koob J, Wolf YI, Koonin EV, Zhang F. 2020. Diverse enzymatic activities mediate antiviral immunity in prokaryotes. Science 369:1077–1084.

28. Rousset F, Depardieu F, Miele S, Dowding J, Laval A-L, Lieberman E, Garry D, Rocha EPC, Bernheim A, Bikard D. 2022. Phages and their satellites encode hotspots of antiviral systems. Cell Host & Microbe 30:740–753.e5.

29. Pinilla-Redondo R, Shehreen S, Marino ND, Fagerlund RD, Brown CM, Sørensen SJ, Fineran PC, Bondy-Denomy J. 2020. Discovery of multiple anti-CRISPRs highlights anti-defense gene clustering in mobile genetic elements. Nat Commun 11:5652.

30. Pawluk A, Bondy-Denomy J, Cheung VHW, Maxwell KL, Davidson AR. 2014. A new group of phage anti-CRISPR genes inhibits the type I-E CRISPR-Cas system of Pseudomonas aeruginosa. mBio 5:e00896.

31. Boyd CM, Angermeyer A, Hays SG, Barth ZK, Patel KM, Seed KD. 2021. Bacteriophage ICP1: A Persistent Predator of Vibrio cholerae. Annu Rev Virol 8:285–304.

32. Brockhurst MA, Harrison E, Hall JPJ, Richards T, McNally A, MacLean C. 2019. The Ecology and Evolution of Pangenomes. Current Biology 29:R1094–R1103.

33. Rubio A, Sprang M, Garzón A, Moreno-Rodriguez A, Pachón-Ibáñez ME, Pachón J, Andrade-Navarro MA, Pérez-Pulido AJ. 2023. Analysis of bacterial pangenomes reduces CRISPR dark matter and reveals strong association between membranome and CRISPR-Cas systems. Science Advances 9:eadd8911.

34. Hamidian M, Nigro SJ. 2019. Emergence, molecular mechanisms and global spread of carbapenem-resistant Acinetobacter baumannii. Microbial Genomics 5:e000306.

35. Tesson F, Hervé A, Mordret E, Touchon M, d’Humières C, Cury J, Bernheim A. 2022. Systematic and quantitative view of the antiviral arsenal of prokaryotes. Nat Commun 13:2561.

36. Starikova EV, Tikhonova PO, Prianichnikov NA, Rands CM, Zdobnov EM, Ilina EN, Govorun VM. 2020. Phigaro: high-throughput prophage sequence annotation. Bioinformatics 36:3882–3884.

37. Camacho C, Coulouris G, Avagyan V, Ma N, Papadopoulos J, Bealer K, Madden TL. 2009. BLAST+: architecture and applications. BMC Bioinformatics 10:421.

38. Rubio A, Garzón A, Moreno-Rodriguez A, Pérez-Pulido AJ. 2023. Biological warfare between two bacterial viruses in a genomic archipelago sheds light on the spread of CRISPR-Cas systems. bioRxiv 10.1101/2023.09.20.558655 (accepted in Cell Reports).

39. Katoh K, Standley DM. 2013. MAFFT multiple sequence alignment software version 7: improvements in performance and usability. Mol Biol Evol 30:772–780.

40. Steenwyk JL, Iii TJB, Li Y, Shen X-X, Rokas A. 2020. ClipKIT: A multiple sequence alignment trimming software for accurate phylogenomic inference. PLOS Biology 18:e3001007.

41. Minh BQ, Schmidt HA, Chernomor O, Schrempf D, Woodhams MD, von Haeseler A, Lanfear R. 2020. IQ-TREE 2: New Models and Efficient Methods for Phylogenetic Inference in the Genomic Era. Mol Biol Evol 37:1530–1534.

42. Xu S, Li L, Luo X, Chen M, Tang W, Zhan L, Dai Z, Lam TT, Guan Y, Yu G. 2022. Ggtree: A serialized data object for visualization of a phylogenetic tree and annotation data. iMeta 1:e56.

43. Couvin D, Bernheim A, Toffano-Nioche C, Touchon M, Michalik J, Néron B, Rocha EPC, Vergnaud G, Gautheret D, Pourcel C. 2018. CRISPRCasFinder, an update of CRISRFinder, includes a portable version, enhanced performance and integrates search for Cas proteins. Nucleic Acids Res 46:W246–W251.

44. Georjon H, Bernheim A. 2023. The highly diverse antiphage defence systems of bacteria. Nat Rev Microbiol 10.1038/s41579-023-00934-x.

45. Tesson F, Planel R, Egorov AA, Georjon H, Vaysset H, Brancotte B, Néron B, Mordret E, Atkinson GC, Bernheim A, Cury J. 2024. A Comprehensive Resource for Exploring Antiphage Defense: DefenseFinder Webservice,Wiki and Databases. Peer Community Journal 4.

46. Yan Y, Zheng J, Zhang X, Yin Y. 2023. dbAPIS: a database of anti-prokaryotic immune system genes. Nucleic Acids Res gkad932.

47. Kogay R, Wolf YI, Koonin EV. 2024. Defence systems and horizontal gene transfer in bacteria. Environmental Microbiology 26:e16630.

48. Wu Y, Garushyants SK, Van Den Hurk A, Aparicio-Maldonado C, Kushwaha SK, King CM, Ou Y, Todeschini TC, Clokie MRJ, Millard AD, Gençay YE, Koonin EV, Nobrega FL. 2024. Bacterial defense systems exhibit synergistic anti-phage activity. Cell Host & Microbe S1931312824000192.

49. Imperi F, Antunes LCS, Blom J, Villa L, Iacono M, Visca P, Carattoli A. 2011. The genomics of Acinetobacter baumannii: insights into genome plasticity, antimicrobial resistance and pathogenicity. IUBMB Life 63:1068–1074.

50. Maestri A, Pons BJ, Pursey E, Chong CE, Gandon S, Custodio R, Olina A, Agapov A, Chisnall MAW, Grasso A, Paterson S, Szczelkun MD, Baker KS, van Houte S, Chevallereau A, Westra ER. 2024. The bacterial defense system MADS interacts with CRISPR-Cas to limit phage infection and escape. Cell Host & Microbe 10.1016/j.chom.2024.07.005.

51. Birkholz N, Jackson SA, Fagerlund RD, Fineran PC. 2022. A mobile restriction-modification system provides phage defence and resolves an epigenetic conflict with an antagonistic endonuclease. Nucleic Acids Res 50:3348–3361.

52. Asakura Y, Kojima H, Kobayashi I. 2011. Evolutionary genome engineering using a restriction-modification system. Nucleic Acids Res 39:9034–9046.

53. Handa N, Nakayama Y, Sadykov M, Kobayashi I. 2001. Experimental genome evolution: large-scale genome rearrangements associated with resistance to replacement of a chromosomal restriction-modification gene complex. Mol Microbiol 40:932–940.

54. Korona R, Levin BR. 1993. Phage-mediated selection and the evolution and maintenance of restriction-modification. Evolution 47:556–575.

55. Yirmiya E, Leavitt A, Lu A, Ragucci AE, Avraham C, Osterman I, Garb J, Antine SP, Mooney SE, Hobbs SJ, Kranzusch PJ, Amitai G, Sorek R. 2024. Phages overcome bacterial immunity via diverse anti-defence proteins. Nature 625:352–359.

56. Asnicar F, Thomas AM, Passerini A, Waldron L, Segata N. 2023. Machine learning for microbiologists. Nat Rev Microbiol 1–15.

57. Kortright KE, Chan BK, Koff JL, Turner PE. 2019. Phage Therapy: A Renewed Approach to Combat Antibiotic-Resistant Bacteria. Cell Host Microbe 25:219–232.

58. Piel D, Bruto M, Labreuche Y, Blanquart F, Goudenège D, Barcia-Cruz R, Chenivesse S, Le Panse S, James A, Dubert J, Petton B, Lieberman E, Wegner KM, Hussain FA, Kauffman KM, Polz MF, Bikard D, Gandon S, Rocha EPC, Le Roux F. 2022. Phage-host coevolution in natural populations. Nat Microbiol 7:1075–1086.

59. Gaborieau B, Vaysset H, Tesson F, Charachon I, Dib N, Bernier J, Dequidt T, Georjon H, Clermont O, Hersen P, Debarbieux L, Ricard J-D, Denamur E, Bernheim A. 2024. Prediction of strain level phage–host interactions across the Escherichia genus using only genomic information. Nat Microbiol 9:2847–2861.

