## Supplementary figures and images for "Innate antiviral systems are major defensome components that drive prophage distribution in *Acinetobacter baumannii*"

### Suppl. Fig. S1

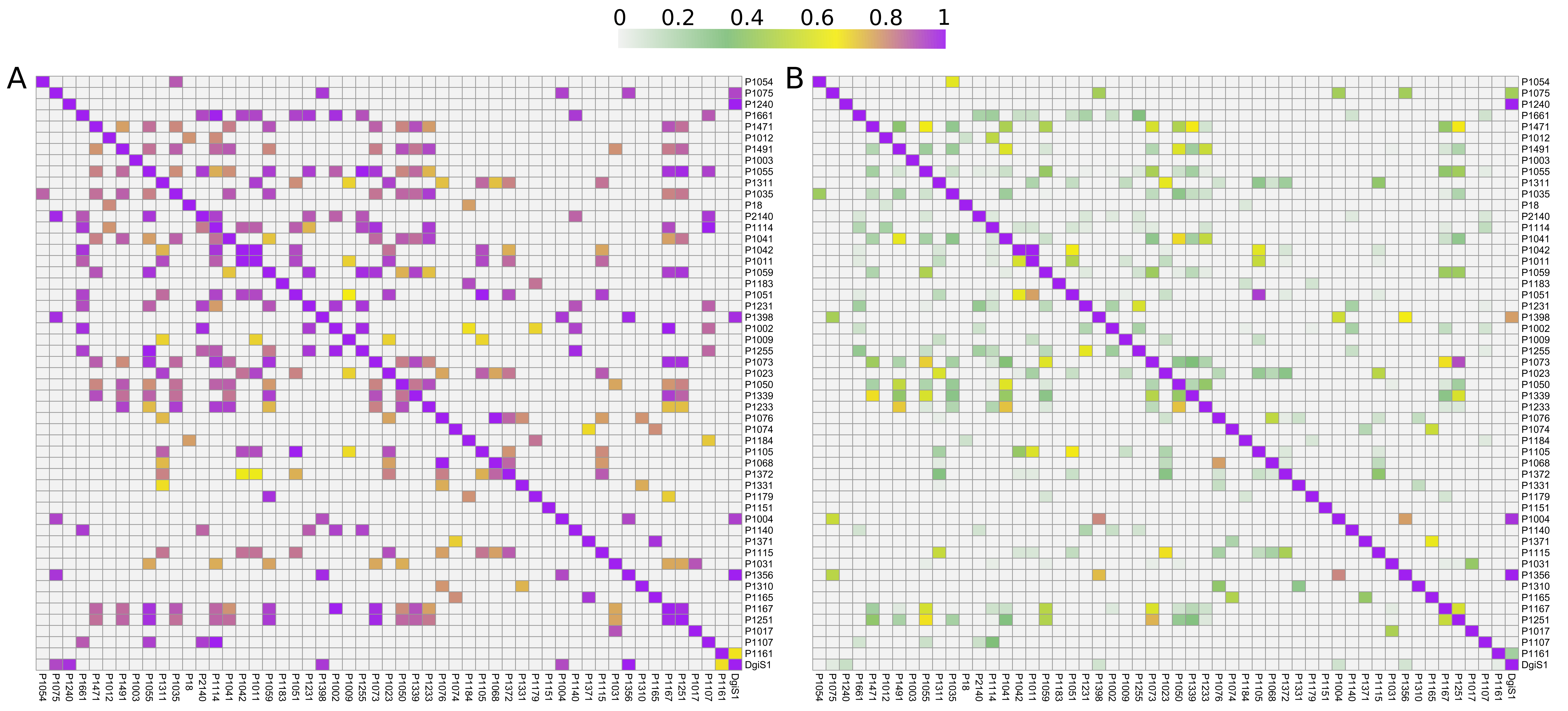

### Suppl. Fig. S2

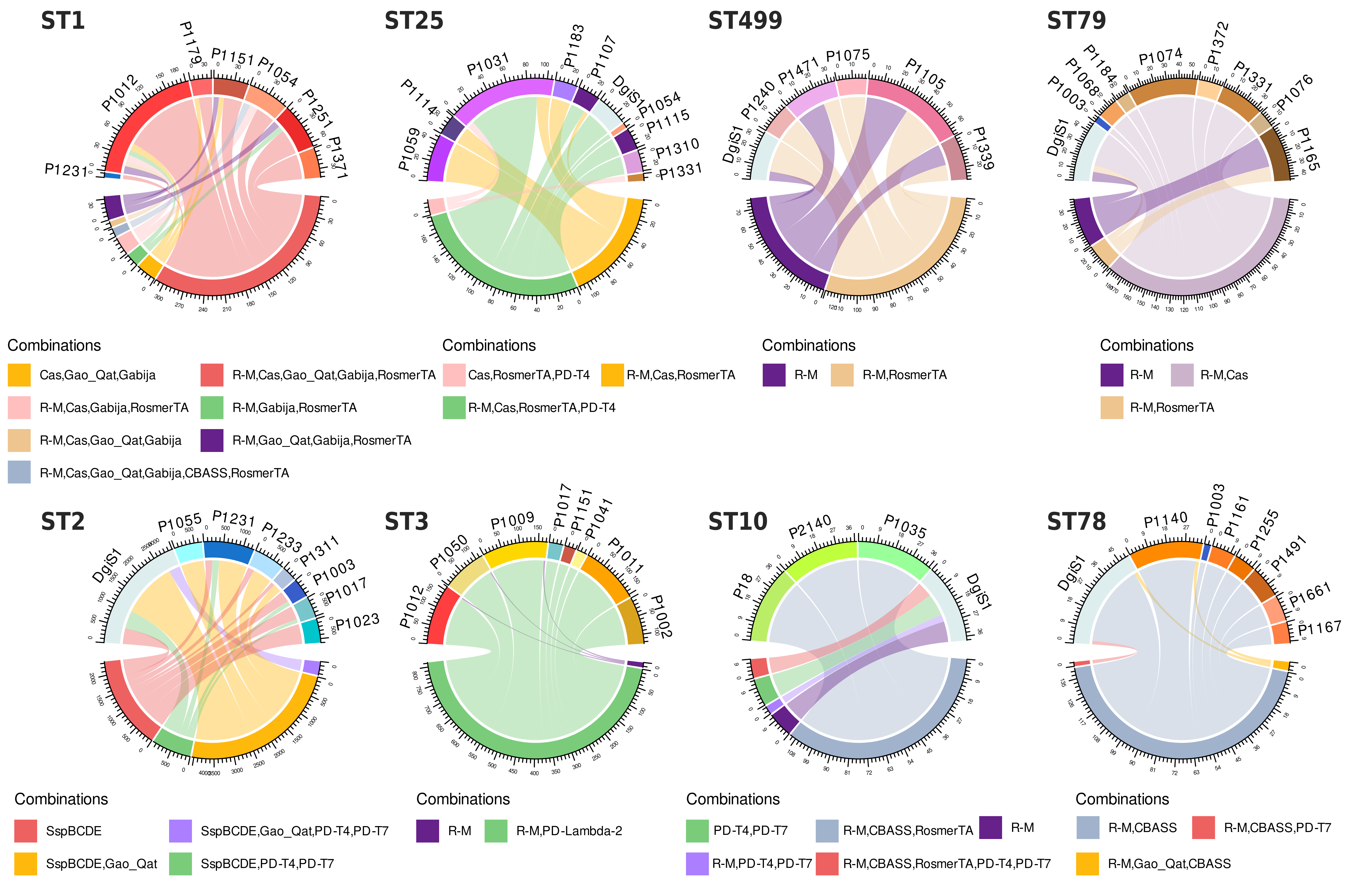
